# Prediction error induced motor contagions in human behaviors

**DOI:** 10.1101/214056

**Authors:** Tsuyoshi Ikegami, Gowrishankar Ganesh, Tatsuya Takeuchi, Hiroki Nakamoto

## Abstract

Motor contagions refer to implicit effects on one’s actions induced by observation of other’s actions. Motor contagions are believed to be induced simply by action observation and cause an observer’s action to become similar to the observed action. In contrast, here we report a new motor contagion that is induced only when the observation is accompanied by *prediction errors*-differences between actions one observes and those he/she predicts or expects. Moreover, this contagion may not manifest as a similarity between one’s own and observed actions. In our experiment, observation of the same action induced distinct motor contagions, depending on whether prediction errors are present or not. In the absence of prediction errors, similarly to previous reports, participants’ actions changed to become similar to the observed action, while in the presence of prediction errors, their actions changed to diverge away from it. Our results suggest distinct effects of action observation and action prediction on human actions.

## Introduction

Our motor behaviors are shaped not just by physical interactions^1–3^ but also by a variety of perceptual^4–8^ interactions with other individuals. *Motor contagions* are the result of such a perceptual interaction. They refer to implicit changes in one’s actions caused by the observation of the other’s actions^9,10^. Studies over the past two decades have isolated various motor contagions in human behaviors, from the so called automatic imitation ^4,11^ and emulation^10,12,13^, to outcome mimicry^14^ and motor mimicry^5,15^. These motor contagions are induced simply by action observation and have a signature characteristic - they cause certain features of one’s action (like kinematics^4,11,16^, goal^10,12,13^, or outcome^14^) to become similar to that of the observed action. In contrast, here we report a new motor contagion that is induced not simply by action observation, but when the observation is accompanied by *prediction errors-* differences between actions one observes and those he/she predicts or expects. Furthermore, this contagion may not lead to similarities between observed actions and one’s own actions. Our results show that distinct motor contagions are induced by the observation of the same actions depending on whether prediction errors are present or not.

## Results

Thirty varsity baseball players participated in our study. The sample size was determined by a power analysis (See Methods). The participants were randomly assigned to one of three groups (n=10 in each): No prediction error (nPE) group, Prediction error (PE) group, and Control (CON) group. The participant’s baseball experience was balanced across the three groups (F(2,27)=1.431, p=0.257, η_p_^2^=0.096). The participants in the nPE and PE groups performed five *throwing* sessions (Fig. 1A) that were interspersed with four *observation* sessions (Figs. 1B, C). The participants in the CON group performed only the throwing sessions. Instead of the observation sessions, they took a break in between the throwing sessions for a time period equivalent to the length of the observation sessions.

**Figure 1:**
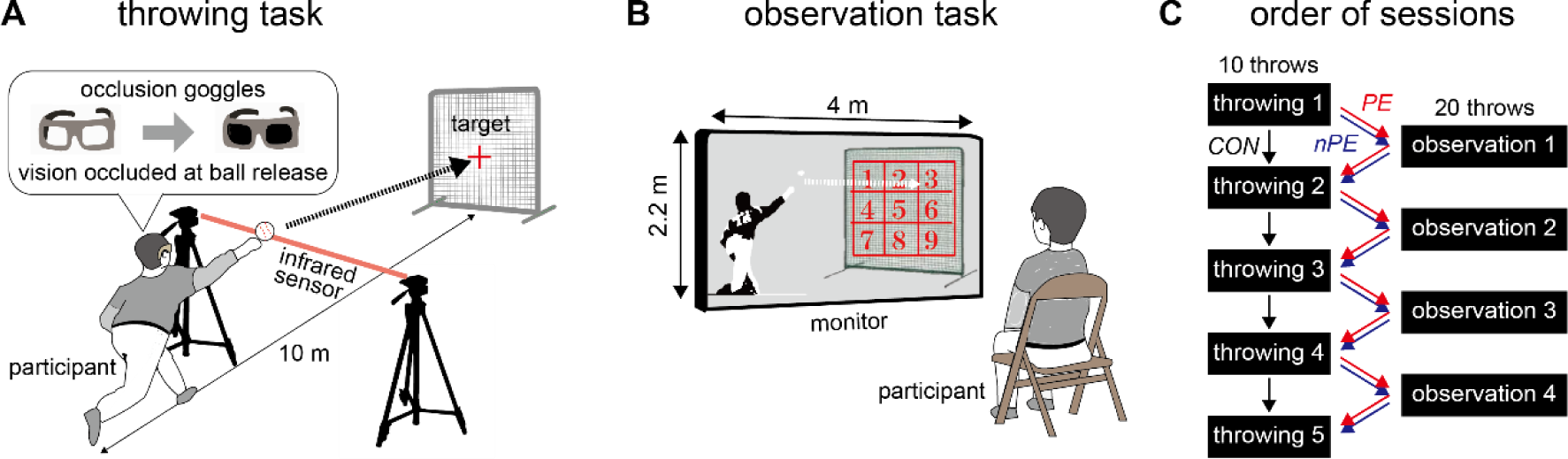
Our experiment consisted of two tasks-A) In the *throwing task* the participants threw balls aimed for the center of a target while wearing “occlusion” goggles, that turned opaque when the participants arm crosses the infrared beam. This prevented them from seeing where their throw hits the target. B) In the *observation task,* the participants were asked to observe the video of throws made by a baseball pitcher and vocally report a number on a nine part grid corresponding to where the throw hit the target. C) Five throwing sessions were interspersed with four observation sessions for the participants in the nPE group (blue arrow) and PE group (red arrow), while the participants in the CON group (black arrow) performed only the throwing sessions, but took a break (equivalent to the length of an observation session) between the throwing sessions.

In the throwing session, the participants in all the groups threw a baseball aiming for the center of a ‘strike-zone’ sized square target placed over the ‘home plate’ (Fig. 1A). They made their throws while wearing “occlusion” goggles that turned opaque when the participants released the ball from their hand (see Fig. 1A). The participants could thus see the target for aiming, but could not see where their ball hits the target. Each throwing session included ten throws.

In the observation session, the participants in the nPE and PE groups watched a video of throws made by an unknown baseball pitcher. After each throw, a numbered grid (see Fig. 1B) appeared on the target (in the video) once the ball hit the target, and the participants were asked to report the grid number corresponding to where they saw the ball hit the target. The purpose of this reporting task was to ensure that the participants maintain their attention on the target in the video. In order to cancel out any spatial bias with respect to the observed actions, half the participants in each group (called upper-right observing participants) were shown throws that predominantly hit the upper right corner of the target (most frequent at #3, see yellow gradient Fig. 2B and methods for observed throw distribution). The other 5 participants (called lower-left observing participants) were shown a video of throws predominantly hitting the lower left corner of the target (most frequent at #7, see Methods). Each observation session included twenty observed throws.

**Figure 2:**
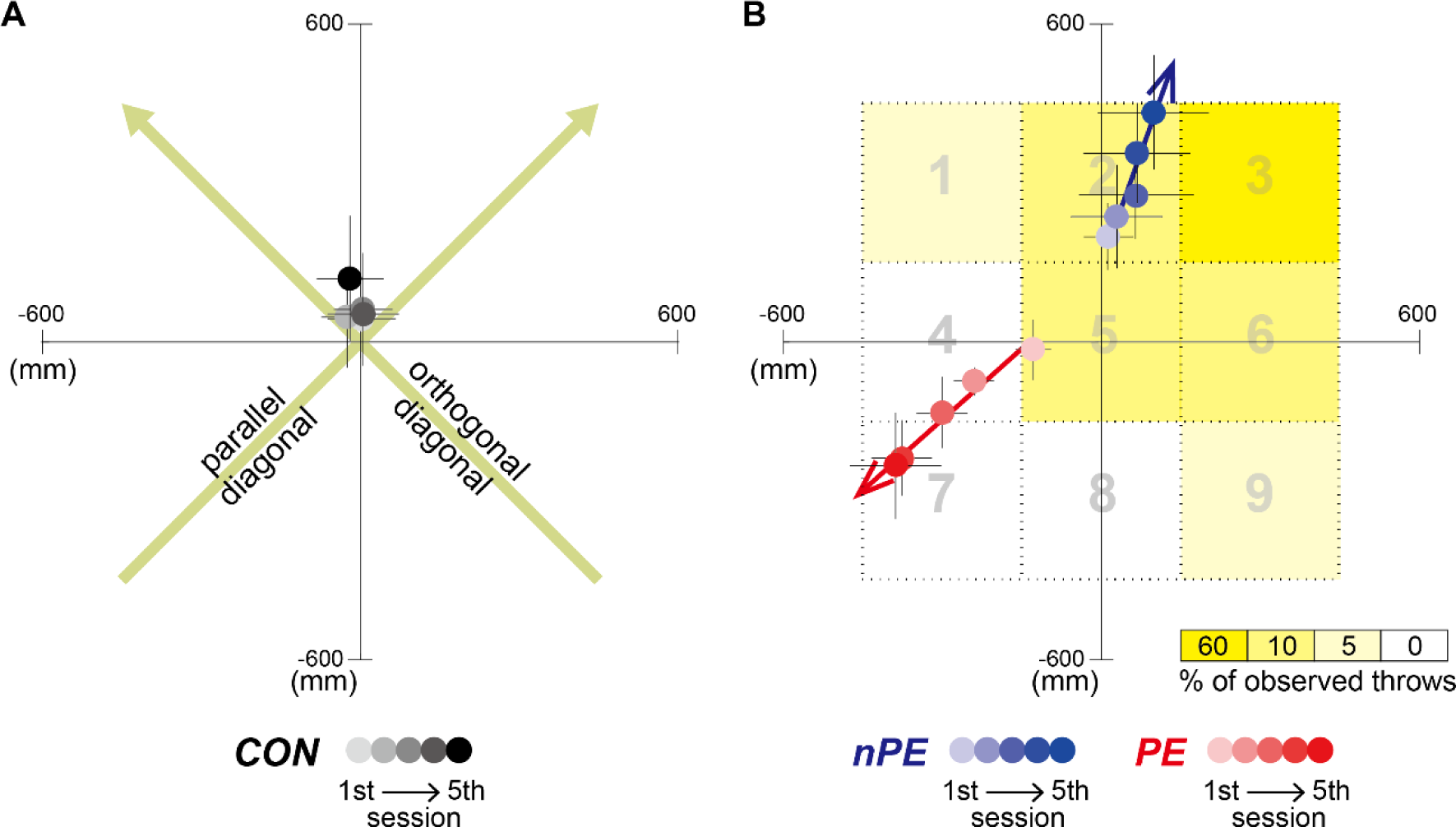
Throw hit locations of participants. A) The across participant average (and s.e.) of hit locations in the CON group are shown in gradients of gray, a darker color representing a later throwing session. The green arrows show the ‘parallel’ and ‘orthogonal’ diagonals, which are used as the reference coordinates for the data plotting and quantification analysis for the nPE and PE groups. B) The across participant average (and s.e.) of hit locations in the nPE group (blue data) and the PE group (red data). The grid area markings shown to the participants after each observed throw, are shown in grey. The yellow gradient indicates the percentage of the observed throws hitting each grid area in the observation sessions for the *upper-right* observing participants, who were shown the pitcher’s throws biased to the upper-right corner of the target. Note that the hit locations were plotted by flipping the data from the *lower-left* observing participants along the parallel and orthogonal diagonals, such that the predominant direction of pitcher’s throws observed by all participants in both the PE and nPE groups were toward the *upper-right* corner of the target. The nPE groups tend to progressively deviate towards to the observed throws, while the PE groups tend to deviate away from the observed throws. The colored arrows indicate the linear fit of the participant-averaged hit locations across the throwing sessions.

Different instructions were provided to the nPE and PE groups in order to manipulate the prediction errors induced in them. The participants in the nPE group were told that “the pitcher in the video is aiming for different grid numbers on the target across trials. These numbers were provided by the experimenter and we display only those trials in which he was successful in hitting the number he aimed for”. On the other hand, participants in the PE group were instructed that “the pitcher in the video is aiming for the center of the target”.

The two different instructions were designed to lead to greater prediction errors in the PE group compared to the nPE group. The instruction to the nPE group prevents the participants from having any prior expectation of the outcome of an observed throw, and was thus expected to attenuate any prediction error. In contrast, the instruction to the PE group makes the participants expect the observed throws to hit near the target center. Similar to previous studies^6,17^, this instruction was thus expected to induce a difference between the throw outcome expected by the participants and the actual outcome observed by them. Specifically, we expected the upper-right and lower-left observation participants in the PE group to experience prediction errors, directed towards the upper right and lower left, respectively.

The participants’ task performance in the throwing session was evaluated as a change in the throw hit location. The position of the hit locations was measured along the ‘parallel’ and ‘orthogonal’ diagonals (see green arrows in Fig. 2A). The diagonal joining the upper-right and lower-left corners, that were the predominant locations of the pitcher’s throws in the observed video, was named as the ‘parallel’ diagonal. Data from the upper-right and lower-left observing participants was analyzed together by flipping the coordinate of the data from the lower-left observing participants (see Methods).

First, in the observation session, the accuracy (% correct) of the report was comparable between the two groups (nPE: 96.25 ± 1.48 (mean±s.d.) %, PE: 95.63 ± 3.32 %; two sample t-test, t(18)=0.516, p=0.612). This ensures that the level of attention to the video was similar between the two groups.

The performance in the throwing session, however, dramatically differed between the two groups. The throws by the nPE group progressively drifted towards where the pitcher in the video predominantly threw the ball (blue data in Fig. 2B). This pattern is similar to the motor contagion reported previously as outcome mimicry^14^. In contrast, the throws by the PE group progressively drifted *away* from where the pitcher in the video predominantly threw the ball (red data in Fig. 2B). The hit locations along the parallel diagonal (Fig. 3) showed a significant interaction between the sessions and groups (F(8,108)=5.124, p=2×10^−5^, η_p_^2^=0.275). Across the sessions, the throws in the nPE group (blue data in Figs. 2B, 3) significantly drifted towards the direction of observed pitcher’s throws (F(4,36)=2.910, p=0.035, η_p_ =0.244; 1 vs 5 sessions by Turkey’s test: p=0.034) while the throws in the PE group (red data in Figs. 2B, 3) significantly drifted away from the observed pitcher’s throws (F(4,36)=5.170, p=2×10^−3^, η_p_^2^=0.365; 1^st^ vs 5^th^ sessions by Turkey’s test: p=5×10^−3^). The throws in the CON group (black data in Figs. 2A, 3) did not show such a drift (F(4,36)=0.297, p=0.878, η_p_^2^=0.032) along the parallel diagonal.

**Figure 3:**
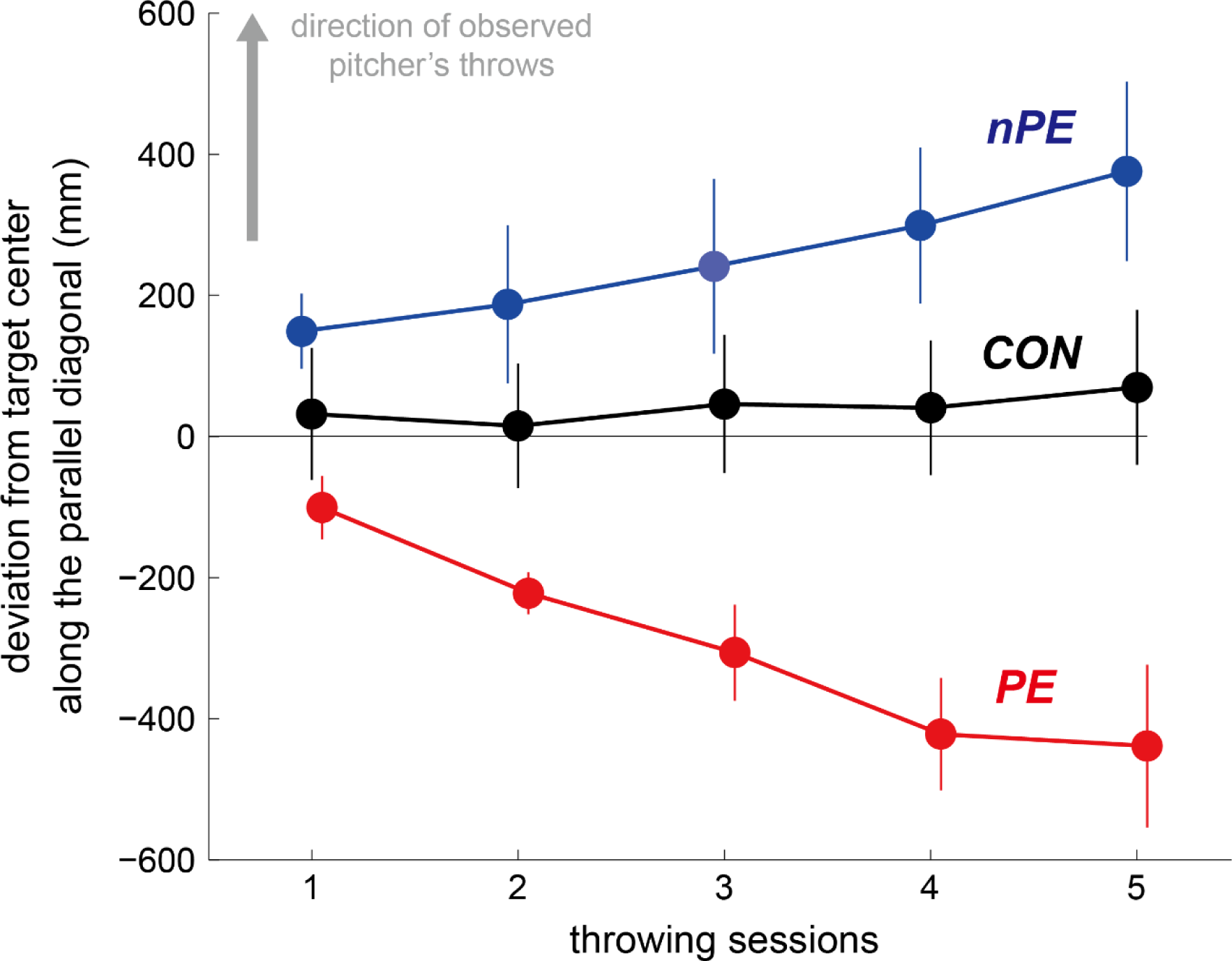
Changes in the hit locations of the participants’ throws across the throwing sessions. The participant-averaged deviations from the target center along the parallel diagonal were plotted by flipping the data from the lower-left observing participants, such that the directions of pitcher’s throws observed by all participants in both the nPE and PE groups were in the positive ordinate (indicated by gray arrow). The data from the CON, nPE, and PE groups are plotted in black, blue and red, respectively. The throws by the nPE group progressively deviated towards where the pitcher in the video threw the ball most frequently. In contrast, the throws by the nPE group progressively deviated away from where the pitcher in the video threw the ball most frequently. The throws by the CON group did not change across sessions. Error bars indicate standard error.

On the other hand, the hit locations along the orthogonal diagonal in all the nPE, PE, and CON groups showed no significant differences across the throwing sessions (F(4,108)=1.762, p=0.142, η_p_^2^=0.061) and between the groups (F(2,27)=1.150, p=0.332, η_p_^2^=0.079). This result confirms that the drifts observed in the nPE and PE groups are not due to possible cognitive fatigue induced by the extra observation task they performed (compared to the CON group), or cognitive biases induced by the validity of the instructions given to them. In either case, we would have expected their throws to also drift in directions other than along the parallel diagonal. The focused drifts along the parallel diagonal suggest that the drifts were induced by the bias in the observed pitcher’s throws (in the nPE group) and the prediction errors (in the PE group), both of which were present specifically along the parallel diagonal.

Together, our results clearly show that the observation of a same action can lead to distinct motor contagions depending on whether the observation takes place in the presence or absence of prediction errors.

## Discussion

Previous motor contagion studies have extensively examined how observation of other’s actions affects action production^4,5,10–12,14,16^. On the other hand, while previous action prediction studies have examined how action production^18,19^, or the ability to produce an action^20–22^ affects prediction of observed actions, the converse, the effect of action prediction on one’s actions, has been rarely examined. One exception is our previous study^6^ which reported that an improvement in the ability to predict observed actions affects the observers’ ability to produce the same action. The affects observed in this study was likely a prediction error induced motor contagion, but as the observed actions and the corresponding prediction errors were not controlled in that study, it is difficult to conclude this cause. Overall, the two effects, of action prediction and action observation on action production, have never been compared. This study compares the two effects for the first time by modulating the prediction errors perceived by participants, when they observe throws by another individual. We show that, while in the absence of prediction errors (nPE group), the action observation makes the participants’ throws to become similar to the observed throws (like in previous reports^4,9,14^), in the presence of prediction errors (PE group), the same action observation makes their throws diverge away from the observed throws (Figs. 2B, 3). Our results thus suggest the presence of a distinct, prediction error induced motor contagion in human behaviors.

The previous motor contagions are believed to be caused by the presence of ‘long term’ sensorimotor associations^4,23^ due to which an observed action automatically activates an individual’s sensorimotor representations of the same action, making his/her actions similar to the observed actions. On the other hand, further studies are required to understand the mechanisms underlying the prediction error induced motor contagion. It is however still interesting to note the parallel between this contagion, and the behavioral characteristic of motor learning. When learning a new motor task, individuals are believed to utilize prediction errors from self-generated actions to change their sensorimotor associations, and correct their actions^24^. The motor contagion in the PE group seems to cause participants to make similar action corrections to compensate for their prediction errors, strangely even though the prediction errors are from actions made by another (observed) individual, rather than from self-generated actions. Therefore, while the previously reported motor contagions may be explained as a facilitation/interference in action production by the sensorimotor associations^4,23^ activated by observed actions, the new contagion we present here is probably a result of erroneous changes in the sensorimotor associations, caused by the observed prediction errors. This mechanism is probably also related to the phenomena of observational motor learning, in which the observation of an action is reported to aid subsequent motor learning^25–28^. This however, remains uncertain because, unlike observational motor learning^29,30^, the contagion effects we observe here are clearly implicit (not willed by the participants), and furthermore, are induced even when the participants are not learning a new motor task, and in fact are arguably over-trained in it.

In conclusion, our results clearly show that action observation and action prediction can induce distinct effects on an observer’s actions. Understanding the interactions between these distinct effects are arguably critical for the complete understanding of human skill learning. For example, it can enable a better understanding of the link between the mechanisms of motor contagions and motor learning, and help develop better procedures to improve motor performances in sports and rehabilitation.

## Acknowledgments

We thank Mr. Kenta Yamamoto for his assistance with the experiments. We thank Dr. Masaya Hirashima and Dr. Nobuhiro Hagura for their valuable comments on this manuscript. This work was partially supported by JSPS KAKENHI Grant #16H05916 (to T.I.), #16K12999 and #26702025 (to H.N.), and #15616710 and #13380602 (to G.G.). The authors declare no competing financial interests.

## Methods

## Participants

Thirty right handed varsity baseball players (20.33±1.62 (mean±s.d.) years old) with normal or corrected vision, took part in this study. Their years of experience in playing baseball was 11.43±1.98 (mean±s.d.). They were randomly assigned to one of three groups of ten participants each: the No prediction error (nPE) group, the Prediction error (PE) group, and the Control (CON) group. All participants were informed of the experimental procedures in advance and consented to take part in this experiment. This study was approved by the local ethics committee in accordance with the Declaration of Helsinki.

The sample size was determined by a power analysis using G*power^31^ (Repeated measure ANOVA within-between interaction, α=0.05, β=0.80, η_p_ =0.06 (medium value)). The power analysis provides the sample size of n=9 for each group. We determined our sample size of n=10 as a more conservative choice with respect to a type-1 error.

## Task and apparatus

All experiments were performed in an indoor baseball facility. The participants in the nPE and PE groups performed two alternating sessions: five throwing and four observation sessions (Fig. 1C), while the participants in the CON group performed only five throwing sessions.

## Sessions

### Throwing sessions

In the throwing sessions, participants in all groups were required to throw a ball aimed at the center of a target placed on the ‘home plate’ 10 m away. The target was a ‘strike zone’ sized square (0.9 m×0.9 m) and its center was indicated by a cross (Fig. 1A). They wore liquid-crystal shutter goggles (PLATO, Translucent Technologies, Toronto) during the throwing sessions. The goggles allowed the participants to see the target when taking aim but occluded their vision for 3 s, starting immediately after they released the ball form their hand in each throw. The timing of ball release as detected by an infrared transmitter-sensor system (AO-S1, Applied Office, Tokyo), that sends a TTL signal to the goggles when the participants’ hand intersects the infrared beam between the transmitter and sensor (Fig. 1A). The position of the infrared transmitter-sensor setup was adjusted for each participant such that the goggle was shut at the timing of their ball release. This system ensured that the participants do not see their ball flight trajectory and the hit location on the target, preventing them from using visual feedback to correct subsequent throws. Each throwing session included ten throws. The ball hit locations were recorded by a digital video camera at 60 fps (Sony HDR-CX560).

### Observation sessions

In the observation sessions, the participants (in the nPE and PE groups) sat on a chair 15 m from a large monitor (2.2 m × 4.0 m) and watched a life-size video of a right-handed baseball pitcher throwing a ball. Unknown to the participants, the pitcher in the video was asked to aim for various areas on the target indicated by an experimenter (see next subsection on observation session videos), and we pre-selected videos to each participant. For five participants in each group, the spatial distribution of throws by the pitcher in the videos were biased to the upper right corner of the target (see Fig. 2B): twelve balls (60 %) in #3; two balls (10 %) in each of #2, #5, #6; one ball (5 %) in each of #1, #9. For the other five participants in each group, the distribution of throws in the videos were biased to the lower left corner of the target (see right panel in Fig. 2B): twelve balls in #7; two balls in each of #4, #5, #8; one ball in each of #1, #9.

Each observation session included twenty video clips and lasted about three minutes. Nine numbered grid areas (0.3 m×0.3 m for each grid) appeared on the target immediately after each throw hit the target and then the participants were asked to vocally announce the grid number corresponding to where the throw hit the target. The participants’ answers were recorded by an experimenter. The task helped us ensure that the participants maintained concentration in the task and watched each throw.

The prediction errors during the observation session were manipulated by a difference of instructions between the nPE and PE groups, provided before each observation session. The participants in the PE group were told that “the pitcher in the video is aiming for the center of the target”. This instruction was expected to generate prediction errors between what the participants expect and what they see in the video. On the other hand, the participants in the nPE group were told that “the pitcher in the video is aiming for various grid numbers on targets. These numbers were provided by the experimenter and we display only those trials in which he was successful in hitting the number he aimed for”. This instruction was expected to prevent the participants from expecting a throw outcome, and hence attenuate prediction errors. The participants in the CON group did not watch a video (they did not have observation sessions) but instead sat on the chair and took a break for three minutes, equivalent to the length of the observation session. The behavior of the CON group was used to check for possible drifts in the participants’ throws due to the persistent lack of feedback in the throwing task.

### Observation session videos

In order to create the video clips for the observation sessions, we recorded movies of a pitcher throwing balls toward the target 10 m away. We utilized a video camera (Sony HDR-CX560, recoding at 60 fps). The camera was placed diagonally (relative to the pitcher–target line) behind the pitcher, 3 m distance away, and recorded the pitcher’s kinematics.

The pitcher was asked to aim his throws to each of the nine areas (#1 to #9) on the target (shown in Fig 1B). He continued to aim for the same target area until he hit it ten times. This procedure provided us with video clips of ten successful throws to each of the nine target areas. From these clips, we chose two clips for throws to #2, #5, #6, (or #4, #5, #8) and one clips for a throw to #1 and #9, and six clips each to throws to #3 (or #7) based on the visibility of ball release and hit location. The six clips (to #3 and #7) were used twice in each observation session.

Next, we edited the selected video clips with Adobe Premiere Pro CS6. Each video clip was temporally clipped from between 2000 ms before ball release to the moment of the target impact. For 3000 ms after the target impact, a grid showing the nine numbered grid areas (Fig. 1B) was superimposed on the target area in the video. The order of presentation of video clips during observation task were randomized among sessions and participants.

## Data analysis

### Hit location analysis

The hit locations recorded by the camera were digitized by using Dartfish (Dartfish, Tokyo, Japan). The hit locations of each throw were first measured in the x-y coordinates where the center of the target was taken as the origin. The throwing task performance in each session were evaluated as the distance of hit location form the target center, averaged over the ten throws for each participant. This value was then averaged across the participants and plotted in Figs. 2 A, B. For statistical analysis, the hit locations by all three groups were analyzed along the diagonals (green arrows in Fig. 2A), parallel and orthogonal to the line joining the corners of the target where the observed throws predominantly landed (#3 or #7). To pool the participants in one group, the hit locations of the throws by the participants who viewed the pitcher aiming for the lower left corner of the target were flipped along the parallel and orthogonal diagonals about the data in the first throwing session. Note that the hit locations in the first throwing session need not be corrected because these represent the default performance by the participant, before the first observations session. We separately performed two-way ANOVAs (3 groups×5 throwing sessions) on the hit locations of the participants’ throws along each of the parallel and orthogonal diagonal. Post hoc pairwise comparisons were performed using the Turkey’s method.

### Observation performance analysis

In the observation session, participants in the nPE and PE groups were required to answer where the balls thrown by the pitchers hit the target by orally reporting one of nine possible areas in each trial. The percentage of correct answers in in each observation session was calculated for each participant.

## References

1 Shergill, S. S., Bays, P. M., Frith, C. D. & Wolpert, D. M. Two eyes for an eye: the neuroscience of force escalation. Science 301, 187, doi:10.1126/science.1085327301/5630/187 [pii] (2003).

2 Ganesh, G. et al. Two is better than one: Physical interactions improve motor performance in humans. Scientific reports 4, doi:ARTN 382410.1038/srep03824 (2014).

3 Takagi, A., Ganesh, G., Yoshioka, T., Kawato, M. & Burdet, E. Physically interacting individuals estimate the partner’s goal to enhance their movements. Nature Human Behaviour 1, 0054, doi:10.1038/s41562-017-0054 https://www.nature.com/articles/s41562-017-0054#supplementary-information (2017).

4 Heyes, C. Automatic imitation. Psychol Bull 137, 463–483, doi:10.1037/a0022288 (2011).

5 Chartrand, T. L. & Bargh, J. A. The chameleon effect: the perception-behavior link and social interaction. J Pers Soc Psychol 76, 893–910 (1999).

6 Ikegami, T. & Ganesh, G. Watching novice action degrades expert motor performance: Causation between action production and outcome prediction of observed actions by humans. Scientific reports 4, 6989, doi:10.1038/srep06989 (2014).

7 Cook, R., Bird, G., Lunser, G., Huck, S. & Heyes, C. Automatic imitation in a strategic context: players of rock-paper-scissors imitate opponents’ gestures. Proc Biol Sci 279, 780–786, doi:10.1098/rspb.2011.1024 (2012).

8 Ganesh, G. & Ikegami, T. in Dance Notations and Robot Motion (eds Jean-Paul Laumond & Naoko Abe) 139–167 (Springer International Publishing, 2016).

9 Blakemore, S. J. & Frith, C. The role of motor contagion in the prediction of action. Neuropsychologia 43, 260–267, doi:10.1016/j.neuropsychologia.2004.11.012 (2005).

10 Becchio, C., Pierno, A., Mari, M., Lusher, D. & Castiello, U. Motor contagion from gaze: the case of autism. Brain 130, 2401–2411, doi:10.1093/brain/awm171 (2007).

11 Brass, M., Bekkering, H. & Prinz, W. Movement observation affects movement execution in a simple response task. Acta Psychol (Amst) 106, 3–22 (2001).

12 Edwards, M. G., Humphreys, G. W. & Castiello, U. Motor facilitation following action observation: a behavioural study in prehensile action. Brain Cogn 53, 495–502 (2003).

13 Gleissner, B., Meltzoff, A. N. & Bekkering, H. Children’s coding of human action: cognitive factors influencing imitation in 3-year-olds. Developmental science 3, 405–414, doi:10.1111/1467-7687.00135 (2000).

14 Gray, R. & Beilock, S. L. Hitting is contagious: experience and action induction. Journal of experimental psychology. Applied 17, 49–59, doi:10.1037/a0022846 (2011).

15 Chartrand, T. L. & van Baaren, R. in Advances in Experimental Social Psychology Vol. Volume 41 219–274 (Academic Press, 2009).

16 Kilner, J. M., Paulignan, Y. & Blakemore, S. J. An interference effect of observed biological movement on action. Curr Biol 13, 522–525 (2003).

17 Ondobaka, S., de Lange, F. P., Wittmann, M., Frith, C. D. & Bekkering, H. Interplay Between Conceptual Expectations and Movement Predictions Underlies Action Understanding. Cereb Cortex, doi:10.1093/cercor/bhu056 (2014).

18 Mulligan, D., Lohse, K. R. & Hodges, N. J. An action-incongruent secondary task modulates prediction accuracy in experienced performers: evidence for motor simulation. Psychol Res, doi:10.1007/s00426-015-0672-y (2015).

19 Hamilton, A., Wolpert, D. & Frith, U. Your own action influences how you perceive another person’s action. Curr Biol 14, 493–498, doi:10.1016/j.cub.2004.03.007 S0960982204001575 [pii] (2004).

20 Knoblich, G. & Flach, R. Predicting the effects of actions: interactions of perception and action. Psychol Sci 12, 467–472 (2001).

21 Urgesi, C., Savonitto, M. M., Fabbro, F. & Aglioti, S. M. Long- and short-term plastic modeling of action prediction abilities in volleyball. Psychol Res 76, 542–560, doi:10.1007/s00426-011-0383-y (2012).

22 Kanakogi, Y. & Itakura, S. Developmental correspondence between action prediction and motor ability in early infancy. Nat Commun 2, 341, doi:ncomms1342 [pii] 10.1038/ncomms1342 (2011).

23 Cook, R., Bird, G., Catmur, C., Press, C. & Heyes, C. Mirror neurons: from origin to function. Behav Brain Sci 37, 177–192, doi:10.1017/S0140525X13000903 (2014).

24 Shadmehr, R., Smith, M. A. & Krakauer, J. W. Error correction, sensory prediction, and adaptation in motor control. Annu Rev Neurosci 33, 89–108, doi:10.1146/annurev-neuro-060909-153135 (2010).

25 Adams, J. A. Use of the model’s knowledge of results to increase the observer’s performance. Journal of Human Movement Studies 12, 89–98 (1986).

26 Schmidt, R. A. & Lee, T. D. Motor control and learning: a behavioral emphasis. 4th edn, (Human Kinetics, 2005).

27 Mattar, A. A. & Gribble, P. L. Motor learning by observing. Neuron 46, 153–160, doi:10.1016/j.neuron.2005.02.009 (2005).

28 Buckingham, G., Wong, J. D., Tang, M., Gribble, P. L. & Goodale, M. A. Observing object lifting errors modulates cortico-spinal excitability and improves object lifting performance. Cortex 50, 115–124, doi:10.1016/j.cortex.2013.07.004 (2014).

29 Maslovat, D., Hodges, N. J., Krigolson, O. E. & Handy, T. C. Observational practice benefits are limited to perceptual improvements in the acquisition of a novel coordination skill. Exp Brain Res 204, 119–130, doi:10.1007/s00221-010-2302-7 (2010).

30 Ong, N. T., Larssen, B. C. & Hodges, N. J. In the absence of physical practice, observation and imagery do not result in updating of internal models for aiming. Exp Brain Res 218, 9–19, doi:10.1007/s00221-011-2996-1 (2012).

31 Faul, F., Erdfelder, E., Lang, A. G. & Buchner, A. G*Power 3: a flexible statistical power analysis program for the social, behavioral, and biomedical sciences. Behav Res Methods 39, 175–191 (2007).

